# Repeated evolution of asymmetric genitalia and right-sided mating behavior in the *Drosophila nannoptera* species group

**DOI:** 10.1101/553024

**Authors:** Andrea Acurio, Flor T. Rhebergen, Sarah Paulus, Virginie Courtier-Orgogozo, Michael Lang

## Abstract

**Background:** Male genitals have repeatedly evolved left-right asymmetries, and the causes of such evolution remain unclear. The *Drosophila nannoptera* group contains four species, among which three exhibit left-right asymmetries of distinct genital organs. In the most studied species, *Drosophila pachea*, males display asymmetric genital lobes and they mate right-sided on top of the female. Copulation position of the other species is unknown.

**Results:** To assess whether the evolution of genital asymmetry could be linked to the evolution of one-sided mating, we examined phallus morphology and copulation position in *D. pachea* and closely related species. The phallus was found to be symmetric in all investigated species except *D. pachea*, which display an asymmetric phallus with a right-sided gonopore, and *D. acanthoptera*, which harbor an asymmetrically bent phallus. In all examined species, males were found to position themselves symmetrically on top of the female, except in *D. pachea* and *D. nannoptera*, where males mated right-sided, in distinctive, species-specific positions. In addition, the copulation duration was found to be increased in *nannoptera* group species compared to closely related outgroup species.

**Conclusion:** Our study shows that gains, and possibly losses, of asymmetry in genital morphology and mating position have evolved repeatedly in the *nannoptera* group. Current data does not allow us to conclude whether genital asymmetry has evolved in response to changes in mating position, or vice versa.

## Background

Changes in behavior are thought to play important roles in animal evolution [1–3]. How new behaviors evolve and how they are encoded in the genome is little understood. New behaviors can initiate secondary evolutionary shifts in morphology, physiology or ecology (“behavioral drive”) [1–9], for example when they bring an organism into contact with new environmental factors. Behavior can also prevent evolutionary changes because plasticity in behavior might enable individuals to adjust for changed environmental conditions [10–12]. Other investigations suggest that behavior and morphology are both subject to natural selection and that their responses to changes in the environment are perhaps independent [13, 14], or that behavior could simultaneously impede and drive evolutionary diversification of different characters [12, 15, 16]. So far, it appears that the effects of behavioral changes on the evolution of morphological traits cannot be generalized and that they require case-specific assessments.

The evolution of left-right asymmetric genitalia in insects is a case where morphology was proposed to have evolved in response to changes in mating behavior [17]. Asymmetric genitalia are observed in many species and phylogenetic studies indicate that they have evolved multiple times independently from symmetric ancestors [18, 19]. While most extant insect species copulate with the male being on top of the female abdomen, the ancestral mating position in insects is inferred to be a configuration with the female on top of the male [18, 20, 21]. The extant male-on-top configuration has likely evolved multiple times in insects [20]. Such changes in mating position probably altered the efficiency of male and female genital coupling, and may have led to the evolution of genital asymmetries to optimize the coupling of genitalia [17].

The *nannoptera* species group belongs to the genus *Drosophila* and consists of four described species that feed and breed on rotten pouches of columnar cacti of the genus *Stenocereus* and *Pachycereus* in Northern and Central America [22–24]. These species are particularly interesting to study the evolution of genital asymmetry because distinct genital structures were identified to be asymmetric in three out of the four described species of this group. *D. acanthoptera* males have asymmetric phallus, *D. pachea* males have a pair of asymmetric external lobes with the left lobe being approximately 1.5 times longer than the right lobe [25, 26], and in the sister species *D. wassermani* males have a pair of asymmetric anal plates (cerci) [25]. In contrast, no asymmetries are known in the fourth described species *D. nannopter*a [27]. The four species separated about 3-6 Ma and lineage-specific changes likely led to the distinct and elaborated asymmetries in each species [28]. Interestingly, *D. pachea* mates in a right-sided copulation position where the male rests on top of the female abdomen with its antero-posterior midline shifted about 6-8 degrees to the right side of the female midline [26, 29]. This one-sided mating posture is associated with asymmetric coupling of female and male genitalia during copulation, with the male genital arch being rotated about 6 degrees towards the female’s right side. Apart from our previous investigations of the *D. pachea* copulation position [26, 29], little is known about mating positions in other *Drosophila* species. In Diptera, several mating positions are known and all involve a symmetric alignment of male and female genitalia. Male and female genitalia are usually inversely positioned relative to each other with the dorsal surface of the aedeagus (phallus) contacting the ventral side of the female reproductive tract [30]. *D. melanogaster, D. simulans* and *D. sechellia* were reported to adopt such a symmetric copulation posture, with the male aligned along the female midline [31–33]. A one-sided mating position was generated artificially in *D. melanogaster* by unilateral ablation of a long bristle located on the genital claspers [31]. In any case, no data is currently available regarding mating positions of the closely related species of *D. pachea*.

The observation of a right-sided mating posture and asymmetric male genitalia in *D. pachea* led us to wonder whether morphological asymmetry in the *nannoptera* group species might have evolved in response to the evolution of one-sided mating [17]. We therefore decided to investigate copulation position and aedeagus asymmetry in species closely-related to *D. pachea*, and to reconstruct their most likely evolutionary history.

## Results

### The phallus of *D. pachea* is asymmetric

The shape of the aedagus/phallus of *D. pachea* has not been described previously. We examined the aedeagus of two dissected *D. pachea* males using scanning electron microscopy (SEM) and found that both were strikingly asymmetric (Fig. 1). Aedeagi were strongly bent, dorsally flattened and pointed at the dorsal tip. Their ventral region bore two ventrally pointing asymmetric spurs, one positioned apically, the other sub-apically. The gonopore was positioned dorso-apically on the right side of the aedeagus. The aedeagal parameres broke off during dissection and were not visualized. In order to corroborate the SEM observations, we dissected and examined 10 aedeagi of *D. pachea* males using light microscopy. Apical and subapical spurs, as well as a right-sided gonopore, were consistently observed in all preparations (n=10, Supplementary Fig.1). Our results indicate that the *D. pachea* phallus is directionally asymmetric (Fig. 2b).

**Figure 1:**
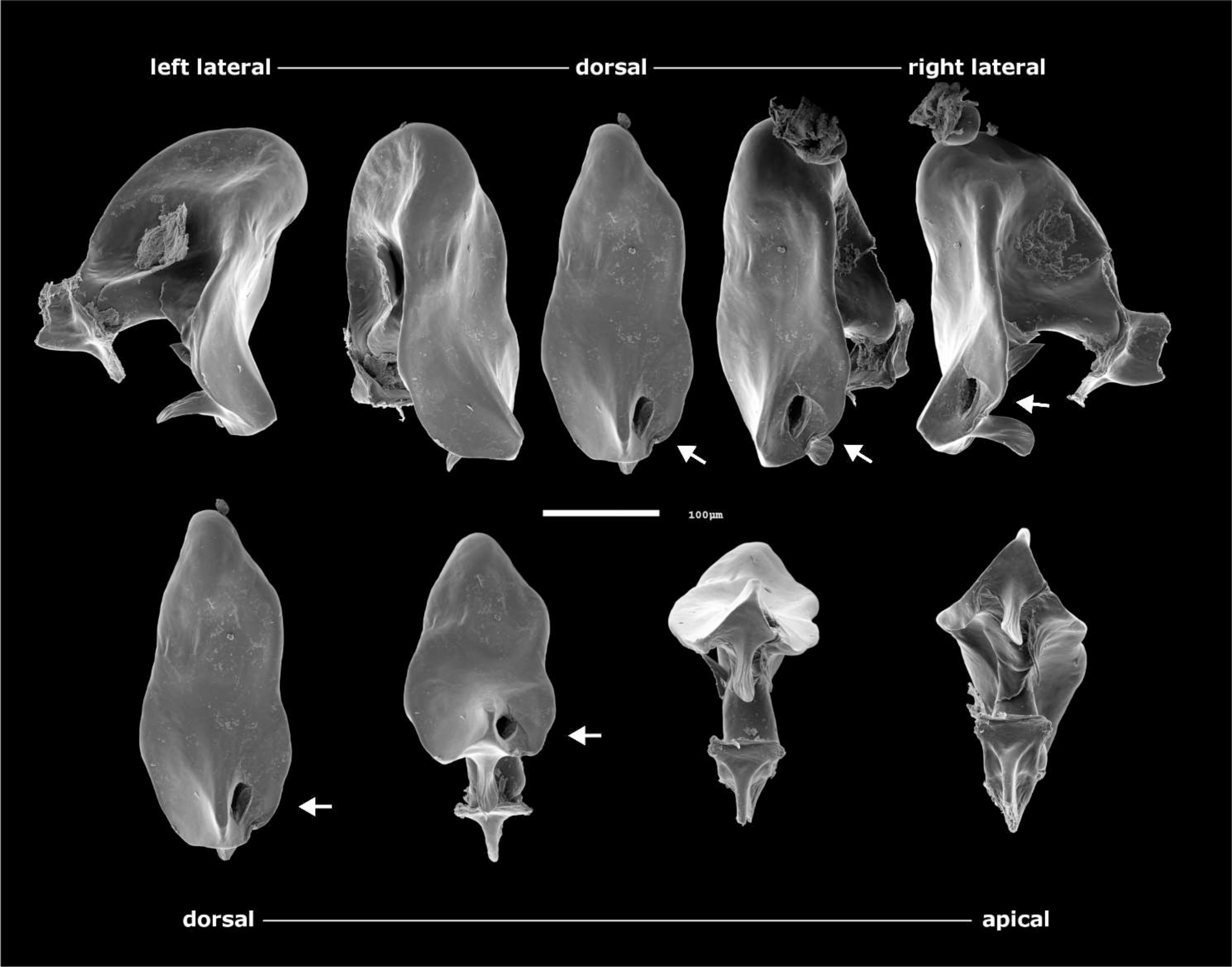
The aedeagus of male *Drosophila pachea* is asymmetric. SEM images of a single phallus in lateral-dorsal and dorsal-apical view. Note the asymmetric position of two subapical spurs, located on the ventral side of the aedeagus, and the asymmetric position of the gonopore. The white arrows point to the gonopore. The scale bar is equivalent to 100 μm.

**Figure 2:**
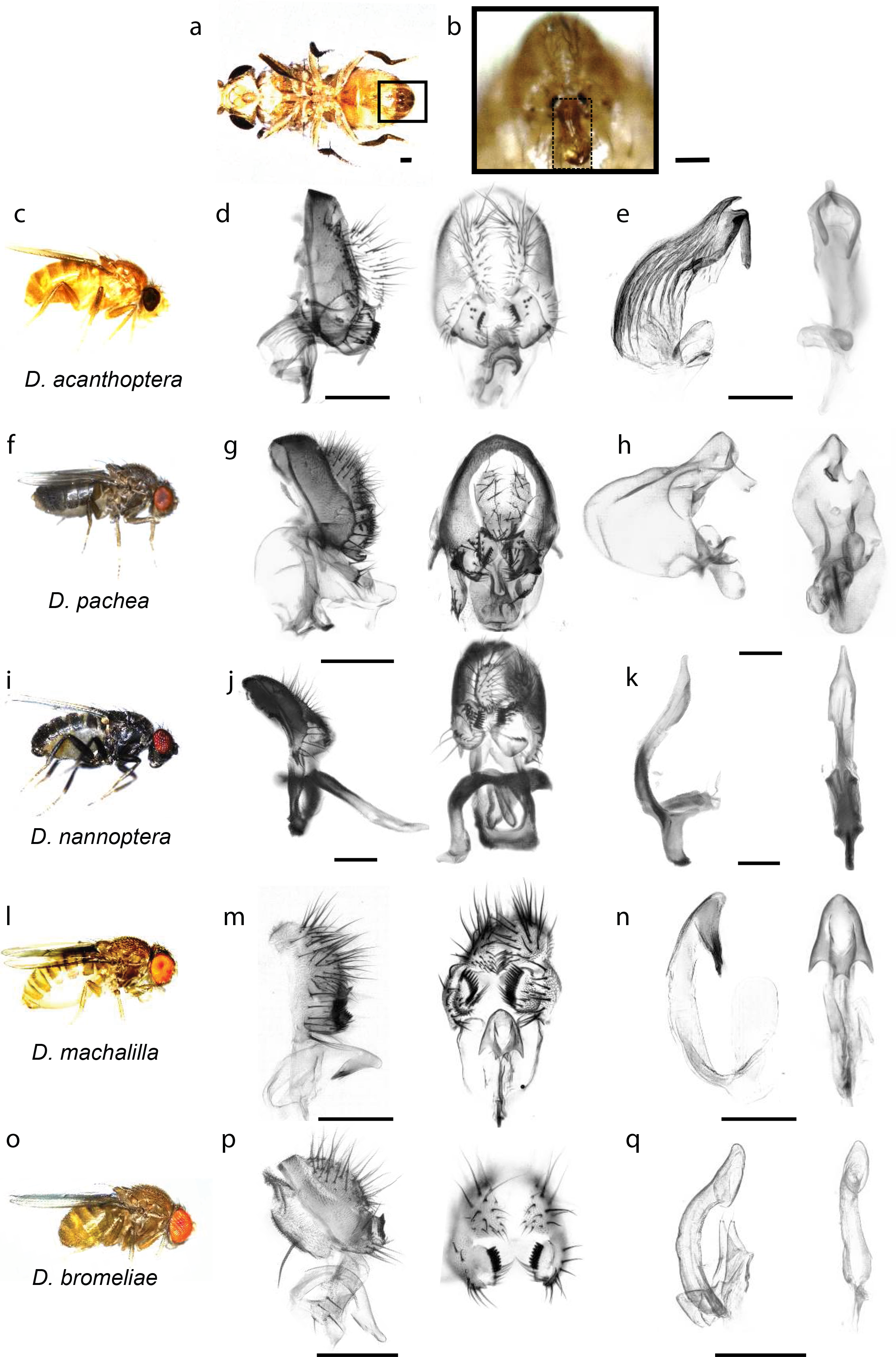
Genital and aedeagus shapes in *D. pachea* and closely related species. External genitalia and aedeagus shapes are compared across closely related species of *D. pachea*. Aedagus asymmetries are only found in *D. acanthoptera* and *D. pachea* (**a)** ventral view of a *D. acanthoptera* male. The black frame indicates the position of male genitalia and the box with a dashed frame shows a magnification with an erected penis. **(b-d, e-g, h-j, k-m, n-p)** Lateral views of male specimen and male genitalia of *D. acanthoptera*, *D. pachea*, *D. nannoptera*, *D. machalilla*, and *D. bromeliae*, respectively. **(c, f, i, l, o)** Male terminalia in lateral and posterior view. **(d, g, j, m, p)** Aedeagus in lateral and ventral view. The scale bar is 100 μm.

### Aedagus asymmetry is observed in *D. acanthoptera* but not in *D. nannoptera*, *D. machalilla* and *D. bromeliae*

We compared aedeagus shapes in several species that are closely related to *D.pachea* (Fig. 2). As previously described [27], the aedeagus of *D. acanthoptera* was found to be asymmetrically bent (n=10). Two asymmetric spurs were found at the ventral apical tip of the aedeagus, with the right spur being consistently longer than the left spur (Fig. 2e, Supplementary Fig.2). However, in contrast to *D. pachea*, no dorso-apical gonopore was observed on the right side of the apex. Aedeagi of *D. nannoptera* males (Fig. 2k, Supplementary Fig. 3) were found to be symmetric (n=15). The ventral side of the apex revealed two apical elongations with slightly variable lengths at the left and right side (n=15, Supplementary Fig.3). The variation in length was not directional and thus considered to reflect random fluctuating asymmetry. The ventral tip of the aedeagus of *D. machalilla (atalaia* species group) (n=10) displayed two lateral hooks (Fig. 2n, Supplementary Fig.4), of the same length on both sides. The aedeagus of *D. bromeliae* showed two lateral symmetric ridges (n=10) (Fig. 2q, Supplementary Fig. 5). In summary, aedeagus asymmetry was only observed in *D. pachea* and *D. acanthoptera*, and distinct phallus structures were found to be asymmetric in these species.

### *D. pachea* and *D. nannoptera* males mate right-sided

The position of the male during copulation has not been described for any of the closely related species of *D. pachea*. In this study, we assessed copulation postures in *D. pachea* and nine related species: *D. acanthoptera* and *D. nannoptera (*sister species of *D. pachea*), *D. machalilla* and *D. bromeliae* (representatives of close outgroup lineages), *D. buzzatii* and *D. mojavensis* (members of the *repleta* species group), as well as representatives of other *Drosophila* species groups (*D. tripunctata*, *D. willistoni* and *D. melanogaster*). Phylogenetic relationships between the ten studied species were estimated with a Bayesian phylogeny (Supplementary Fig. 6) based on a previously published sequence dataset [28], supplemented with publicly available sequence data (this study) for *D. tripunctata* and *D. willistoni*. The obtained phylogeny is congruent with previous findings [28] that *D. nannoptera*, *D. acanthoptera* and *D. pachea* form a monophyletic group with a short internode branch length between the split of the *D. nannoptera* lineage and the separation of *D. acanthoptera* and *D. pachea*. Also, *D. machalilla* and *D. bromeliae* form two close outgroup lineages of the *nannoptera* clade [28, 34, 35], followed by the *repleta* group species *D. buzzatii* and *D. mojavensis* [28].

For each species, we introduced a single virgin female and a single virgin male into a circular mating chamber and recorded the couple until copulation ended or for 45 min when no copulation was detectable. We obtained 315 movies, of which 111 were used for assessing courtship duration, 146 for copulation duration and 124 for copulation posture analysis (supplementary dataset 3). Most movies were discarded because no copulation occurred or individuals had damaged wings or legs (all reasons listed in supplementary dataset 3). As previously described [36–38], copulation duration varied significantly among species (ANOVA, df1 = 9, df2 = 136, F = 73.38, p < 2e-16) (Table 1). We could reproduce a previously reported trend that copulation duration in *nannoptera* group species was remarkably long compared to *D. mojavensis* and *D. buzzatii* of the *repleta* group, with copulation duration of 88.49 min ± 35.18 min for *D. acanthoptera*, 29.58 ± 7.86 min for *D. pachea* and 11.9 ± 4.2 min for *D. nannoptera* (mean ± SD). In comparison, copulation duration of *D. buzzatii* 1.79 ± 0.65 min and *D. mojavensis* 2.3 ± 0.35 min (mean ± SD) of the *repleta* species group was shorter and similar to *D. machalilla* 2.28 ± 0.53 min and *D. bromeliae* 0.92 ± 0.28 min (mean ± SD) (Table 1).

**Table 1:**
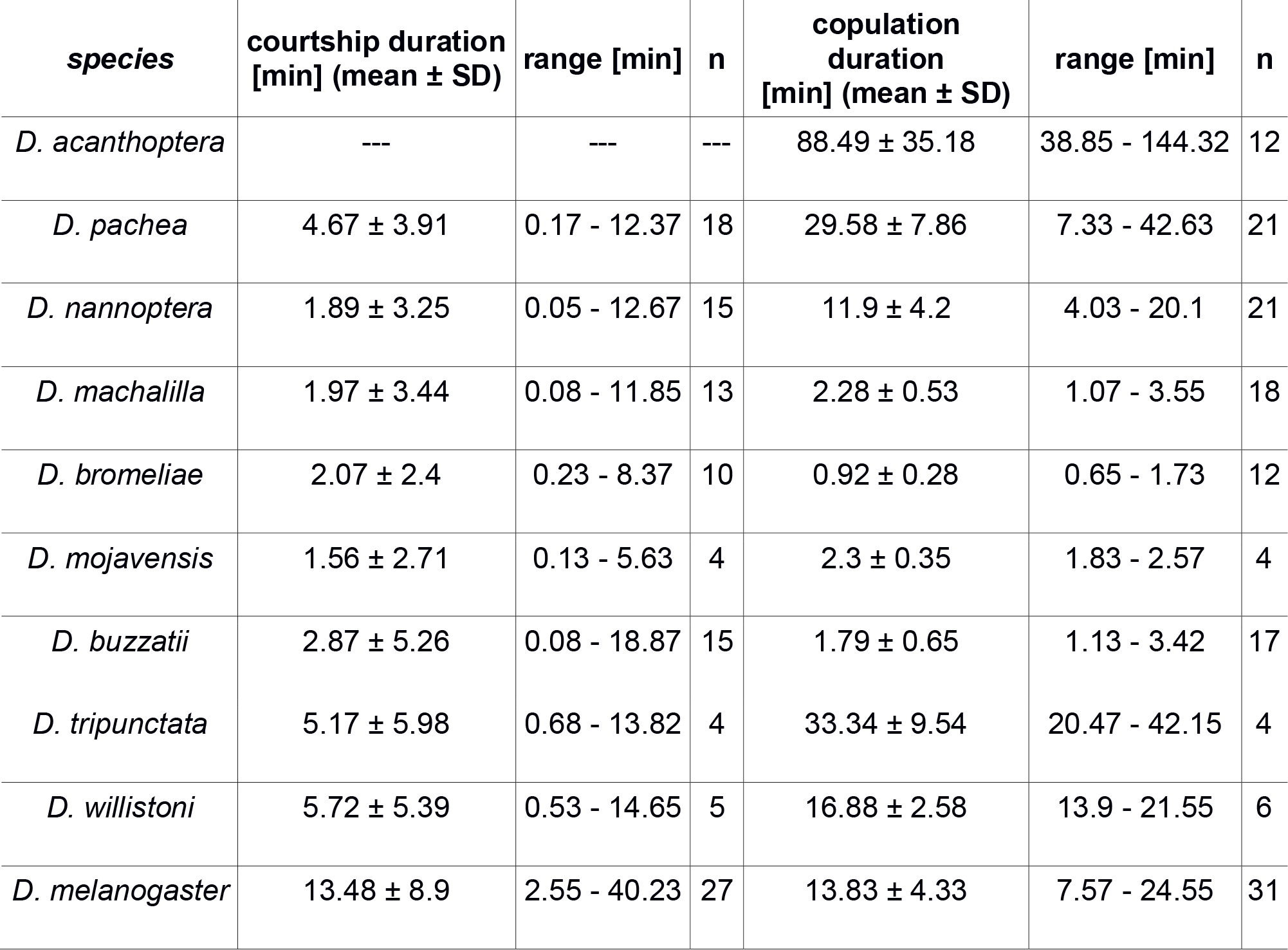
Courtship and copulation duration.

| *species* | courtship duration<br>[min] (mean ± SD) | range [min] | n | copulation<br>duration<br>[min] (mean ± SD) | range [min] | n |
| --- | --- | --- | --- | --- | --- | --- |
| *D. acanthoptera* | --- | --- | --- | 88.49 ± 35.18 | 38.85 - 144.32 | 12 |
| *D. pachea* | 4.67 ± 3.91 | 0.17 - 12.37 | 18 | 29.58 ± 7.86 | 7.33 - 42.63 | 21 |
| *D. nannoptera* | 1.89 ± 3.25 | 0.05 - 12.67 | 15 | 11.9 ± 4.2 | 4.03 - 20.1 | 21 |
| *D. machalilla* | 1.97 ± 3.44 | 0.08 - 11.85 | 13 | 2.28 ± 0.53 | 1.07 - 3.55 | 18 |
| *D. bromeliae* | 2.07 ± 2.4 | 0.23 - 8.37 | 10 | 0.92 ± 0.28 | 0.65 - 1.73 | 12 |
| *D. mojavensis* | 1.56 ± 2.71 | 0.13 - 5.63 | 4 | 2.3 ± 0.35 | 1.83 - 2.57 | 4 |
| *D. buzzatii* | 2.87 ± 5.26 | 0.08 - 18.87 | 15 | 1.79 ± 0.65 | 1.13 - 3.42 | 17 |
| *D. tripunctata* | 5.17 ± 5.98 | 0.68 - 13.82 | 4 | 33.34 ± 9.54 | 20.47 - 42.15 | 4 |
| *D. willistoni* | 5.72 ± 5.39 | 0.53 - 14.65 | 5 | 16.88 ± 2.58 | 13.9 - 21.55 | 6 |
| *D. melanogaster* | 13.48 ± 8.9 | 2.55 - 40.23 | 27 | 13.83 ± 4.33 | 7.57 - 24.55 | 31 |

**Table 2:** Test for one-sided mating positions. Fit: GLM (angle ~ species), family = “gaussian”, hypothesis: angle = 0, Bonferroni corrected p-values

| *species* | settling |  |  | 10% stable copulation time point<br>replicate measurement 1 |  |  | 10% stable copulation time point<br>replicate measurement 2 |  |  |
| --- | --- | --- | --- | --- | --- | --- | --- | --- | --- |
|  | est.<br>contrast | z<br>value | p | est.<br>contrast | z<br>value | p | est.<br>contrast | z value | p |
| *D. acanthoptera* | -1.9105 | -0.282 | 1 | 0.5919 | 0.091 | 1 | -0.4980 | -0.075 | 1 |
| ***D. pachea*** | **21.4011** | **4.048** | **0.000517** | **18.8120** | **3.704** | **0.00212** | **17.8498** | **3.449** | **0.00564** |
| ***D. nannoptera*** | **32.5346** | **6.646** | **3.00e-10** | **34.4231** | **7.321** | **2.46e-12** | **35.0479** | **7.314** | **2.60e-12** |
| *D. machalilla* | 5.7585 | 0.851 | 1 | 6.5023 | 1.001 | 1 | 8.2219 | 1.242 | 1 |
| *D. bromeliae* | -3.9911 | -0.590 | 1 | -2.9702 | -0.457 | 1 | -0.7399 | -0.112 | 1 |
| *D. mojavensis* | 3.0034 | 0.268 | 1 | 4.2370 | 0.393 | 1 | 6.3650 | 0.580 | 1 |
| *D. buzzatii* | 2.4878 | 0.430 | 1 | 3.0853 | 0.555 | 1 | 1.2320 | 0.217 | 1 |
| *D. tripunctata* | 3.6189 | 0.323 | 1 | -2.9770 | -0.239 | 1 | -5.4349 | -0.429 | 1 |
| *D. willistoni* | 0.7288 | 0.080 | 1 | 0.4118 | 0.047 | 1 | 0.6928 | 0.077 | 1 |
| *D. melanogaster* | 6.6156 | 1.414 | 1 | 1.6010 | 0.356 | 1 | 0.7997 | 0.175 | 1 |

To assess mating posture, we calculated the angle between a line drawn through the male head midline and the female scutellum tip and a second line drawn through the female head midline and the female scutellum tip (Supplementary Fig. 7A). The angle was set positive when male head lies on the right side of the female and negative when on the left. The camera view relative to the fly couple position within the mating cell may affect the measured angle in each experiment but the sign of the average mating angle taken from different recordings for each species should accurately reflect the one-sidedness of the male mating position. As a consequence, we expected a one-sided copulation position to produce a consistent positive or negative distribution of angle values, while symmetric mating positions should result in an angle distribution around zero.

To compare mating angles between species, it is necessary to examine copulation postures at the same corresponding time point during copulation. At copulation start, the male position on top of the female was found to be greatly variable between couples, even within a single species, so this time point was not considered appropriate for our comparative analysis. Since copulation duration varies greatly between species, finding another comparable time point across species was not trivial. We subdivided copulation into two phases, an initial phase where the male is on top of the female abdomen but consistently moving legs and abdomen, and a second phase when the male maintains an invariant position relative to the female, which can sometimes walk or move its legs (Supplementary Fig. 6). The “settling time point” is defined as the time point between the first and second phase, when the male adopts an invariant position relative to the female. For our cross-species analysis we chose to assess copulation angle at two time points: (1) right after the male had settled into an initial invariant copulation position (the settling time point) and (2) at 10% of elapsed time between the settling time point and the end of copulation (10% stable copulation time point). For species with a mean copulation duration > 2.5 min, > 15 min or > 60 min, we also measured the angles every 2.5 min, 5 min or 10 min, respectively. This allowed us to follow mating postures of each species over the course of copulation.

Significant one-sided mating positions were observed in *D. pachea* and *D. nannoptera*, both at the settling time point and at the 10% stable copulation time point (Fig.3a,b, Table 2). No significant one-sided copulation postures were detected in *D. acanthoptera* and the other seven tested species including *D. melanogaster* (Fig. 3a,b).

**Figure 3:**
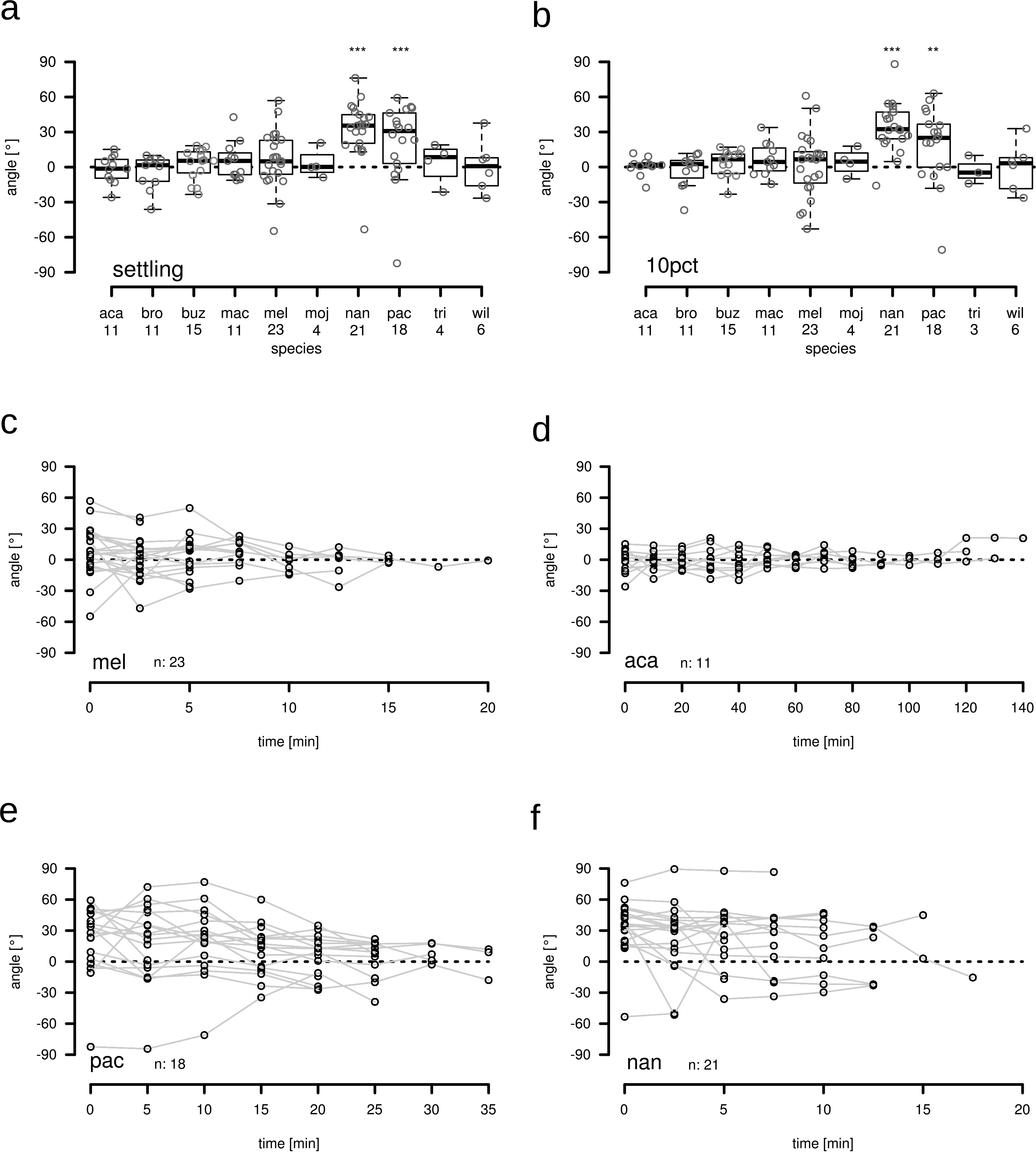
The copulation position of *D. nannoptera* and *D. pachea* is asymmetric. Copulation angles of *D. pachea* couples and of nine related *Drosophila* species; aca: *D. acanthoptera*, bro: *D. bromeliae*, buz: *D. buzzatii*, mac: *D. machalilla*, mel: *D. melanogaster*, moj: *D. mojavensis*, nan: *D. nannoptera*, pac: *D. pachea*, tri: *D. tripunctata*, wil: *D. willistoni*. **(a,b)** Copulation angle at the settling time point (settling, see material and methods) and at the 10% stable copulation time point (10pct), respectively. Stars indicate significant rejection of the null hypothesis: angle = 0 (Table 2, GLM fit angle~species **: p<0.001, ***: p< 0.0001). Numbers below each boxplot indicate the number of observations. The dashed lines indicate an angle of zero degrees. **(c-e)** Copulation angles over the course of copulation of *D. melanogaster* (mel), *D. acanthoptera* (aca), *D. pachea* (pac) and *D. nannoptera* (nan). n indicates the number of observations. Grey lines connect points obtained from the same copulation couple over time. The dashed lines indicate an angle of zero degrees.

Over the course of copulation, mating angles continued to range over zero for *D. melanogaster* and *D. acanthoptera* (Fig. 3c,d), indicating a relatively steady symmetric copulation position without any left- or right-sidedness. Similar to previous investigations [26, 29], *D. pachea* revealed right-sided angles that were highest at the beginning of copulation at 0-10 min after settling (Fig. 3e). At later time points, the angles tended to range over zero. In *D. nannoptera*, mating angles tended to be right-sided throughout copulation (Fig. 3f). In summary, *D. pachea* and *D. nannoptera* revealed a right-sided copulation posture whereas all the other tested species displayed a symmetric mating posture.

### Male *D. nannoptera* tilt to the right side of the female abdomen during copulation

To further investigate the right-sided copulation posture in *D. nannoptera* and better observe the male position relative to the female dorso-ventral midline, we filmed the couples from a frontal perspective (Supplementary Fig. 8). In particular, we assessed the inclination of the male body relative to the female dorso-ventral axis by measuring the angle P4-P5-P6, with P4 as the medial most dorsal edge of the female head (often visible by the ocelli), P5 being the most ventral medial position of the female head (the female proboscis) and P6 as the medial most dorsal edge of the male head (often visible by the ocelli) (Supplementary Fig. 8).

*D. nannoptera* mating positions were on average strikingly right-sided (Supplementary Fig. 8), with a considerable variation of observed angles, ranging from slightly left- to strongly inclined right-sided (−8.42° – 57.7°) over the course of copulation. Left-sided angles were only observed during the first two minutes of copulation. On average, the male tended to initially adopt a right-sided copulation posture with an angle of 10.36° ± 6.88° (mean ± SD) (n=25) between 0-1 min after copulation start (Table 3). Over the course of copulation, the angle then increased to 27.16° ± 10.81° (n=29) between 3-4 min after copulation start (Table 3), which was visible by an inclination of the male head towards the female’s right side. This tilt-movement was not observed in *D. pachea*, where all males remained on top of the female abdomen [29]. We therefore conclude that *D. pachea* and *D. nannoptera* adopt distinct copulation postures, even though both of them are right-sided.

**Table 3:** *D. nannoptera* frontal mating angles. The mean estimates for each time interval were calculated with average values when multiple measurement points were available for a given experiment

| time interval after cop. start [ min] | angle (mean ± SD) | n |
| --- | --- | --- |
| 0 - 1 | 10.36 ± 6.88 | 25 |
| 1 - 2 | 15.46 ± 8.82 | 29 |
| 2 -3 | 23.44 ± 12.15 | 29 |
| 3 - 4 | 27.16 ± 10.81 | 27 |
| 4 - 5 | 29.1 ± 11.62 | 25 |
| 5 - 6 | 28.97 ± 11.92 | 21 |
| 6 - 7 | 32.48 ± 8.29 | 13 |
| 7 - 8 | 26.44 ± 10.24 | 5 |
| 8 - 9 | 23.21 ± 11.88 | 4 |
| 9 - 10 | 24.7 ± 13.66 | 3 |
| 10 - 11 | 18.65 ± 14.03 | 2 |
| 11 - 12 | 8.07 ± NA | 1 |

## Discussion

### Phallus asymmetries differ between *D. pachea* and *D. acanthoptera*

The currently published data suggest that genital asymmetries are rare among *Drosophila* species. The genus *Drosophila* encompasses over 1500 described species [39] and only 8 species have been shown without doubt to display an asymmetric phallus: *D. marieaehelenae* and *D. hollisae* of the flavopilosa group [40, 41], *D. asymmetrica* and *D. quinarensis* of the guarani group [42, 43] *D. endobranchia* of the canalinea group [44], *D. acuminanus* and *D. freilejoni* of the onychophora group [27, 45, 46] and the *nannoptera* group species *D. acanthoptera* [27]. Genital asymmetry might be more widespread than what is reported in the literature across *Drosophila*, as certain species are only described based on a few specimens, and subtle asymmetric characters might have been overlooked and interpreted as fluctuating variation between left and right sides. Here, we compared aedeagus morphology of at least 10 specimens of five different species that belong to the *nannoptera* species group and closely related species. We did not detect aedeagus asymmetry in the tested species outside of the *nannoptera* species group and found that within the *nannoptera* group only *D. acanthoptera* and *D. pachea* but not *D. nannoptera* reveal striking left-right asymmetries (Fig. 4). We did not evaluate aedeagus asymmetry of *D. wassermani*, as this species is not available for examination and our attempts to catch specimen in the wild were not successful (see materials and methods). Asymmetries differed between *D. pachea* and *D. acanthoptera*. Whereas ventral spurs on the *D. pachea* aedeagus were apart from each other, with one being apical and the other subapical, *D acanthoptera* aedeagus had a pair of apical spurs that differed in length. In addition, the gonoopore was visible and right-sided in *D. pachea* while it was not visible in *D. acanthoptera*. Our results thus highlight that the asymmetric phallus structures of *D. pachea* and *D. acanthoptera* are derived morphologies that have little in common and diversified independently after the split of the two species about 3-6 Ma ago [28]. It is impossible to infer whether the asymmetries observed in both species derived from a pre-existing asymmetric phallus in their ancestor or if asymmetry evolved *de novo* in both lineages.

**Figure 4:**
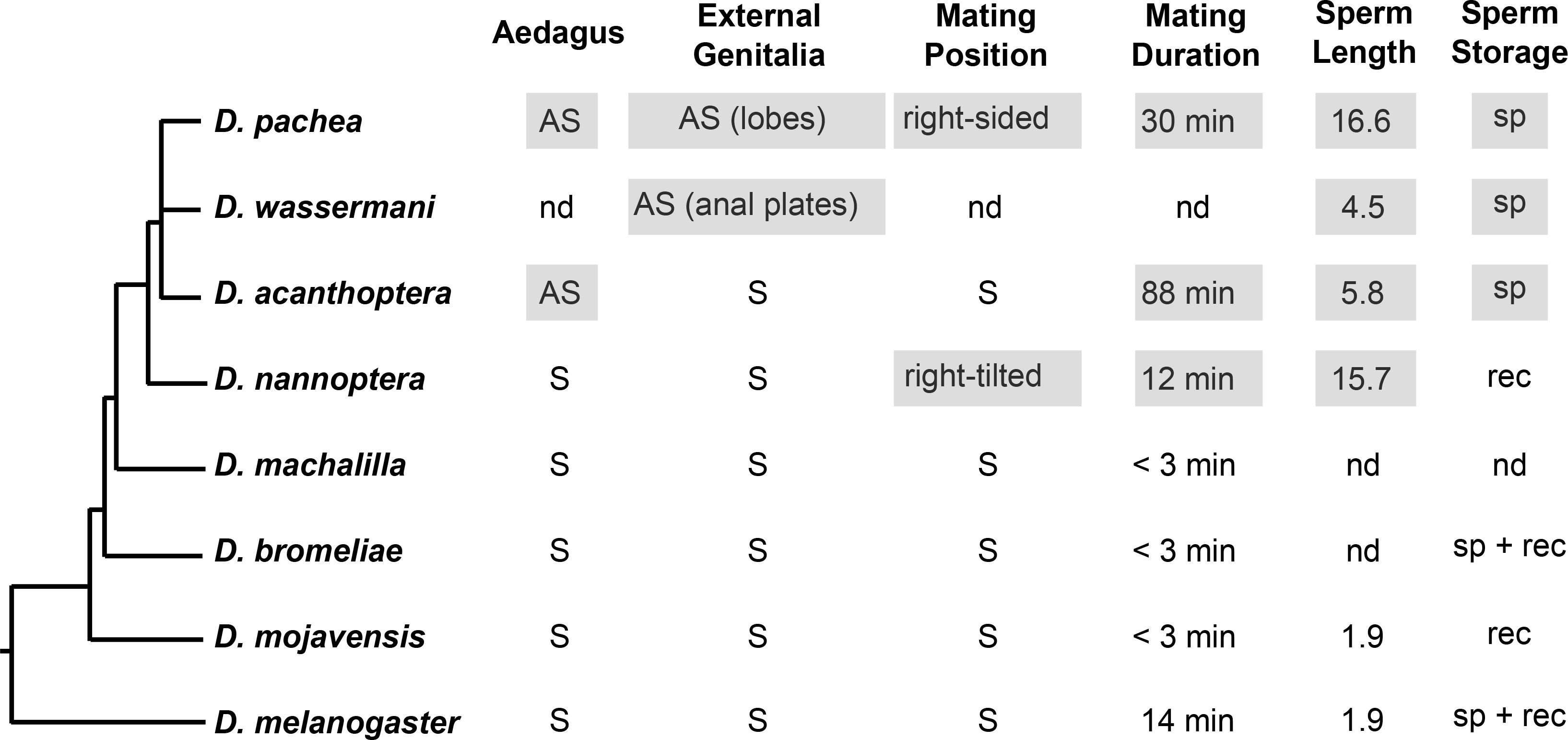
Evolution of sexual characters in the *nannoptera* species group. The cladogram was established based on the phylogeny of this study (supplementary Fig. 6), combined with data from Lang et al (2014) [28]. AS, asymmetric states; S, symmetric states; nd, not determined, sp, spermathecae; rec, female seminal receptacle.

The outer genitalia (epandrium) has been reported to be asymmetrical in *D. pachea* (where the left lobe is longer than the right lobe [25, 26] and in *D. wassermani* (where the right anal plate is larger than the left one [25]). Our inspection of a few dissected epandria of *D. nannoptera*, *D. acanthoptera* and *D. machalilla* revealed no obvious asymmetry (Fig. 2). However, a quantitative comparison remains to be done to confirm the absence of asymmetry in the epandrium of these species.

### Long copulation duration is specific to the nannoptera group species

We observed that *nannoptera* species copulated considerably longer than any representative species of the close outgroup lineages (Fig. 4). This trend was previously reported by Pitnick and Markow (1991) [36, 37] where the authors compared copulation duration of *nannoptera* group species with *repleta* group and other species. Here we included two additional closely related species, *D. machalilla* and *D. bromeliae*, and observed that their copulation durations were relatively short. Our observations therefore indicate that a long copulation duration is specific to the *nannoptera* group.

### Right-sided mating positions differ between *D. pachea* and *D. nannoptera*

We assessed copulation postures of *D. pachea* and a range of related species to track the conservation of right-sided mating position in the *nannoptera* group. Two aspects of copulation behavior made cross-species comparisons difficult. First, copulation duration was extremely diverse and ranged from less than a minute in *D. bromeliae* to more than two hours in *D. acanthoptera*. Second, the movements of the male and female during copulation varied between species. In *D. melanogaster* and *D. willistoni*, we observed vigorous movements of the male during copulation accompanied by female hindleg kicking. These phases were interrupted by periods without movements. In contrast, males of most other species initially moved upon mounting the female and then settled into an invariant copulation posture relative to the female. We chose to compare mating postures across species once the couple adopted the invariant position, at the settling time point, and at 10% of elapsed time between the settling time point and the end of copulation (10% stable copulation time point). These two time points were assumed to represent comparable moments during copulation.

At the two measured time points during copulation, the angle between the male midline and the female midline during copulation was distributed symmetrically around zero, indicating a symmetric mating position in all tested species except *D. pachea* and *D. nannoptera*. Our previous data from *D. pachea* [26, 29] was re-analysed in this study with a different measurement approach and led to the same conclusion as our earlier reports. In addition, we found that *D. nannoptera* adopts a right-sided mating position with angle values that were slightly higher than in *D. pachea* (Fig. 3). Assessment of homology of behaviors is difficult compared to morphological characters, because Owen’s position criterium for homology [47] does not exist for behavioral traits. Observation of similar behaviors does not necessarily mean common descent [6]. Our precise examination of the mating position of *D. nannoptera* from a frontal perspective revealed that *D. nannoptera* males strongly tilt to the female’s right side during copulation, a behavior that is not observed in *D. pachea* [26, 29]. Therefore, mating postures can be regarded as distinct between the two species. Interestingly, a comparable tilting behavior during copulation was observed in experiments with *D. pachea* males that had surgically modified external genital lobes [29]. Male lobes are considered to be important in grasping the female abdomen beneath the oviscapt valves and to keep *D. pachea* upright on the female abdomen. A hypothetical scenario is thus that the ancestral mating position in shared ancestor of the two species may have been right-tilted but the evolution of asymmetric external lobes in *D. pachea* led to a derived right-sided copulation posture, which is upright. Alternatively, right-sided mating position may have evolved independently in the two lineages leading to *D. pachea* and *D. nannoptera*. In all scenarios, at least two evolutionary changes in mating position must be considered to account for the distinct, species-specific right-sided mating positions in the *nannoptera* group.

### Asymmetry in mating position and in phallus have evolved in different branches of the *nannoptera* group phylogeny

Across the *nannoptera* group, we find no striking correspondence between right-sided mating posture and asymmetric male genitalia. For example, *D. acanthoptera* has an asymmetric aedeagus but mates in a symmetric overall posture. On the opposite, no directional asymmetry is detected in the male (external and internal) genitalia of *D. nannoptera*, but males adopt a right-sided copulation posture (Fig. 4). Based on our phylogeny, *D. nannoptera* presents the earliest branching lineage within the *nannoptera* group. In this sense, right-sided mating postures could have originated earlier during evolution than asymmetric morphologies and may have been lost in *D. acanthoptera*. However, the internode branch length between the split of the *D. nannoptera* lineage and the separation of *D. acanthoptera* and *D. pachea* is short and statistical support is weak [28]. Thus, phylogenetic relationships within the *nannoptera* group remain to be resolved and it is more appropriate to regard all *nannoptera* species as sister species.

So far, we conclude that both right-sided copulation behavior and asymmetric male genitalia evolved within the *nannoptera* species group and that diversification of both traits have involved lineage-specific evolutionary changes. They may have evolved by modifications of pre-existing right-sided mating behavior and/or asymmetric genital morphologies already present in the ancestor. Alternatively, they can have appeared *de novo* in each extant lineage.

### One-sided mating and asymmetric phallus are correlated with giant sperm and female sperm storage, respectively

Asymmetric genital morphology and right-sided mating behavior may also be associated with other characters that are special to the *nannoptera* species group. *D. pachea* and *D. nannoptera* are among the *Drosophila* species that produce the longest (giant) sperm [48, 49] (Fig. 4). The association of right-sided mating with giant sperm production actually holds better than with asymmetric male genital morphology because *D. acanthoptera* has an asymmetric aedeagus but has relatively small sperm [48] and mates in a symmetric overall posture (Fig. 4). A specific one-sided mating posture might be necessary for optimal transfer of giant sperm. Examining mating postures in *Drosophila* species which harbor even longer sperm (*D. bifurca* 58 mm, *D. kanekoi* 24 mm, *D. hydei* 23 mm, *D. eohydei* 18 mm) [50, 51] would be interesting to test further the possible association between sperm length and one-sided mating.

The species *D. pachea*, *D. acanthoptera* and *D. wassermani* are also special in the way the female stores sperm after copulation (Fig. 4). They are the only *Drosophila* species that store sperm exclusively inside the spermathecae but not in the seminal receptacle as most other species [49]. In contrast, *D. nannoptera* stores sperm exclusively inside the seminal receptacle [49]. Morphological phallus asymmetry is thus observed in those species that reveal exclusive sperm storage in the spermathecae. Male specimen of *D. wassermani* are thus required to analyse their phallus shapes to confirm this trend. On the other hand, it is hard to generalize from our observations as only three species are concerned.

## Conclusion

Phallus asymmetries were identified in *D. pachea* and *D. acanthoptera* of the *nannoptera* species group and distinct structures were observed to be asymmetric in both species. An increased copulation duration was found to be specific to *nannoptera* group species and was not observed in the closely related outgroup species *D. machalilla* and *D. bromeliae*. Right-sided mating positions were detected in *D. pachea* and *D. nannoptera* and were found to be distinct between them. Our data does not allow us to conclude whether the evolution of the right-sided copulation position may have promoted the evolution of genital asymmetry, or vice versa. Our results nevertheless indicate that asymmetry in genital morphology and in copulation behavior have evolved through multiple evolutionary steps in the *nannoptera* group, revealing a complex history of sexual trait changes, maybe in relationship with the evolution of giant sperm and unique sperm storage in the *nannoptera* group.

## Methods

### Fly sampling and maintenance

An isofemale stock of *Drosophila machalilla* was established from a collection of A. A. in December 2015 at San Jose Beach (01°13′46.4″S, 80°49′14.6″W) Ecuador, using a modified version of the fly traps described in [52]. Our baits contained rotten pieces of the columnar cactus *Armatocereus carwrightianus* and yeast solution. The *D. machallila* stock was raised on standard *Drosophila* medium (60 g/L brewer’s yeast, 66.6 g/L cornmeal, 8.6 g/L agar, 5 g/L methyl-4-hydroxybenzoate and 2.5% ^v^/_v_ ethanol) and a piece of fresh Opuntia *ficus-indica* (prickly pear opuntia) or *Hylocereus undatus* (dragon fruit) in the medium. The isofemale stock was raised for two generations before experiments started and it was maintained for a total of 36 generations.

We also intended to collect *D. wassermani* in August 2016 in Oaxaca, Mexico. Six localities were sampled based on previous records: Reserva de la Biosfera Tehuacan-Cuicatlan (18°11′21.30″ N, 97°14′ 51.7″ W), Huajuapan de Leon (17°48′25.6″ N, 97°14′ 56.7″ W), San Luis del Rio (16°46′30″ N, 96°10′ 49.9″ W), and four sites along the Carretera Internacional 190: Kms 73 (16°42′57.2″ N, 96°19′41.9″ W), 89 (16°40′41.3″ N, 96°14′41.7″ W), 102 (16°42′11.3″ N, 96°11′32.4″ W) and 111 (16°39′48.4″ N, 96°07′31.8″ W). We used banana traps, cactus baits that contained rotten organ pipe cactus *Stenocereus prionosus* and mixed food traps that additionally contained banana and yeast. Besides the invasive species *Zaprionus indianus* and cosmopolitan species *D. melanogaster* and *D. simulans*, we identified several species of the *repleta* group, about 100 individuals of *D. nannoptera*, three males of *D. wassermani* and one female of *D. acanthoptera*. Unfortunately, we were not successful in establishing iso-female strains from *D. nannoptera* and *D. acanthoptera*.

All other stocks were retrieved from the San Diego Drosophila Species Stock Center or were provided by Jean David (Supplementary Table 1). Flies were maintained at 25 °C, except for *D. melanogaster*, *D. tripunctata* and *D. willistoni*, which were either maintaind at 22 °C or 25 °C (see supplementary dataset 3 for details). Flies were kept in vials with 10 mL of standard *Drosophila* food medium (see above) inside incubators with a 12 h light: 12h dark photo-periodic cycle combined with a 30-min linear illumination change between light (1080 lumen) and dark (0 lumen). For maintenance of *D. pachea*, we mixed standard *Drosophila* food medium in the food vial with 40 μL of 5mg/mL 7-dehydrocholesterol (dissolved in ethanol) [53].

### SEM analysis of the *D. pachea* aedeagus

Virgin males of at least 14 days after hatching from the pupa were transferred into a 2 mL reaction tube, snap frozen in liquid nitrogen and stored in ethanol at −20°C. For dissection, frozen and fixed males were placed in 80% ethanol at room temperature and the aedeagus was dissected out with fine needles. Tissues were dried using an EM CPD300 automated critical point dryer (Leica) and mounted on aluminium stubs with the distal end facing upwards and coated with platinum/palladium (20 nm). Each aedeagus was SEM-imaged with a JSM-7500F field emission scanning electron microscope (Jeol) at 270x magnification.

### Analysis of aedeagus asymmetry by light microscopy

The terminal segments of the male abdomen were picked out with fine forceps and boiled for 10 min in two drops of 30% KOH. Genital parts were further dissected on a microscope slide (Thermo Scientific Menzel) in a drop of water using 0.1 mm Minutien Pins (Fine Science Tools) under the stereo-microscope K-500 (VWR). Dissected structures were mounted in pure glycerol on 1.5 mm concave microscope slides (Marienfeld). Images were acquired with a light microscope VHX2000 (Keyence) equipped with a zoom lens VH-Z100UR/W at 350-550 fold magnification. For storage, male genitalia were mounted in DMHF medium on microscope slides (Entomopraxis).

### Phylogenetic analysis

We used data of eight species from a multi-locus dataset of Lang et al. (2014) [28], and added corresponding sequences for *D. willistoni* and *D. tripunctata* (Supplementary Table 2, Supplementary Datafile 2, BEAST input file in DRYAD). For *D. tripunctata*, only mitochondrial data was available at GenBank (https://www.ncbi.nlm.nih.gov/nucleotide/) and missing data was annotated by ‘?’. Phylogenetic analysis was performed in BEAST [54] according to the settings described in Lang et al (2014) [28]. Markov-Chain Monte-Carlo (MCMC) runs were performed with a chain length of 10^7^ generations and recorded every 1000 generations. MCMC output analysis was carried out using TreeAnnotator [54] and the tree was visualized and edited with FigTree [55]. We chose a strict molecular clock and set priors for most recent common ancestors according to the divergence estimates of Lang et al. (2014) [28] for the splits of *D. nannoptera* - *D. pachea* 3.7 ± 1.5 Ma, *D. bromeliae* - *D. pachea* 8 ± 3 Ma. The divergence estimate for all analyzed species was set to 40 ± 5 Ma [56].

### Copulation recording

Emerged flies (0-14 h) were anesthetized with CO_2_, separated according to sex and transferred into food vials in groups of either 5 females or 5 males using a Stemi 2000 (Zeiss) stereo microscope and a CO_2_-pad (Inject+Matic sleeper). Flies were maintained at 22°C or 25°C until they reached sexual maturity (Supplementary Table 1). Males of *D. bromeliae*, *D. melanogaster*, *D. pachea* and *D. nannoptera* were isolated into single vials for at least two days before the experiment was performed. For video recording, one male and one female were introduced with a self-made fly aspirator by mouth suction into a circular plastic mating cell with a diameter of 10-12 mm, a depth of 4mm and a transparent 1-mm Plexiglas cover [26]. For copulation recording of *D. acanthoptera*, flies were let to initiate copulation in a food vial and were then rapidly transferred to the mating cell.

Movies were recorded in a climate controlled chamber [26] at 22 or 25 ± 0.1°C and 60% or 85% ± 5% humidity (supplementary datafile 3). Flies were filmed from above using digital microscope cameras 191251-62 (Conrad), DigiMicro Profi (DNT) or MIRAZOOM MZ902 (OWL). Movies were recorded with the program GUVCVIEW (version 0.9.9) GTK UVC or Cheese (version 3.18.1) (https://wiki.gnome.org/Apps/Cheese) at a resolution of 800 × 600 pixels on a Linux Ubuntu operating system. Movies were recorded until copulation ended or for at least 45 min when no copulation was detectable. After movie recording, flies were dissected or stored in ethanol at −20°C.

### Multiple species mating position analysis

Each movie name consisted of a three-letter abbreviation for the species filmed, an additional two-digit number that also indicated the species and a two-digit number for each respective experiment. Movies were analyzed with the video editor OpenShot 1.4.3 (Open Shot Studios, Texas, USA). Courtship start, copulation start, the settling time point and the end of copulation were annotated manually by two different persons, except for movies of *D. pachea* and *D. melanogaster*, which were annotated only by one person (supplementary datafile 3). Courtship was defined to start when the male displayed at least three consecutive typical courtship behaviors, such as tapping the female, following the female’s abdomen, licking the female oviscapt or the ground beneath the female abdomen, wing rowing (*D. melanogaster*) or other wing vibrations [57]. Courtship was defined to end with the start of copulation, when a male started to mount the female abdomen. Only cases where the male remained mounted on the female for at least 15 sec were counted as copulation starts. Copulation was defined to end when the male had completely descended from the female abdomen with the forelegs detached from the female dorsum and female and male genitalia being separated. As mentioned above, the male moved its legs and abdomen for a certain time period (considered as the settling phase) until adopting an invariant abdomen posture at the settling time point (supplementary datafile 3, Supplementary Fig. 6). The remaining copulation period was defined as the stable copulation period (Supplementary Fig. 6). In fact, this period was often interrupted by periods of vigourous movements in *D. melanogaster*, *D. tripuncata* and *D. willistoni*. In the other species, males remained rather invariant on the female abdomen after the settling time point.

We video-recorded 315 movies, of which 111 were used for assessing courtship duration (supplementary dataset 3). Reasons for discarding 204 movies for courtship duration measurements were: wrong handling of the camera or the software, damaged files: 4; incomplete recording of courtship: 43; fly leg or wing damaged: 27; no copulation after 45 min of experiment start: 129; wing damaged and incomplete recording of courtship: 1. A total of 146 movies were used for the analysis of copulation duration. Reasons for discarding 169 movies for copulation duration measurements were: wrong handling of the camera or the software, damaged files: 4, incomplete recording of copulation: 7, fly leg or wing damaged: 27, no copulation after 45 min of experiment start: 129; multiple reasons: 2 (supplementary dataset 3). From these 146 movies, we had to exclude 22 movies for the assessment of the copulation posture because landmark positions could not be observed. This was mainly due to couples being recorded from the ventral view. As a result, 124 experiments were used for assessment of the copulation posture (supplementary dataset 3). One additional movie was discarded for posture assessment at the 10% stable copulation time point because the female head was not in the camera field of view.

Movie names were replaced by a seven-digit random number (supplementary datafile 5) so that mating postures were quantified in a blind fashion with respect to the species name. Time points for position analysis (supplementary dataset 3, supplementary Fig. 6) were calculated with a custom R script and exported values were used as an input for a bash script to extract images from each movie at particular time points with avconv (libav tools, https://www.libav.org).

The angle was measured using three landmarks on the female and male body: the anterior tip of the female head along its mid-line (P1), the distal tip of the female scutellum (P2) and the most posterior medial point of the male head (P3) (supplementary Fig. 7a). In cases where images were too dark, positions of P1 and P3 were approximated as the anterior and posterior mid distances between the eyes and the position of the scutellum tip (P2) was approximated by the medial dorsal point at the body constriction observed between the third thoracic and first abdominal segment. Position landmarks were placed manually on each image using imageJ and data analysis was done using R. Briefly, coordinates (supplementary dataset 4) were rotated and scaled, so that all P1 points were superimposed and all P2 points as well (supplementary Fig. 7B-K). The angle P1-P2-P3 (Supplementary Fig. 7A) was used to measure one-sidedness of mating positions (Fig. 3). Repeatability of landmark positioning was assessed by two independent rounds of coordinate acquisition for all species at one specific time point during copulation, the 10% stable copulation time point (see text) (2x 124 images). Variation in angle estimates was found to be attributable mostly to individual images and not to replicate measurement (ANOVA, linear model: angle ~ image + replicate, image: df1 = 122, df2 = 123, F = 87.174, p < 2e-16, replicate: df1 = 1, df2 = 244, F = 0.077, p = 0.782).

Hypothesis testing was performed in R to compare mating postures across species (Fig. 3) with the null hypothesis: angle = 0, using the functions glm for generalized linear model fits, and glht to derive estimated contrasts.

### Analysis of the *D. nannoptera* copulation posture

Flies were reared and isolated before copulation as described above. One female and one male were CO_2_ anesthetized and transferred onto a white plastic support (mating cap) and were caged with a transparent plastic cylindrical 25 mm × 7 mm cap. Once courtship was observed, mating caps were put on a motorized horizontally turning stage (0-30 rpm) (grinding stone 8215, Dremel) in front of a camera MIRAZOOM MZ902 (OWL) and copulation was recorded with the camera being put into an optimized frontal view towards the female head by rotating or turning the mating cap. The transparent cap was optionally removed once copulation had started. The yield of informative experiments with these settings was poor as we performed 167 mating experiments but only 29 experiments were informative for our data analysis (supplementary datafile 6, reasons for discarding the experiments are listed). Images were extracted with avconv (see above) every 15-30 sec or when the flies were visible in a frontal view. We measured the inclination of the male body relative to the female dorso-ventral axis by using three landmarks: P4 as the medial most dorsal edge of the female head (often visible by the ocelli), P5 being the most ventral medial position of the female head (the female proboscis) and P6 as the medial most dorsal edge of the male head (often visible by the ocelli) and measuring the angle between the lines drawn through P4-P5 and P5-P6 (Supplementary Fig. 8, supplementary datafile 7).

## List of abbreviations

SEM: scanning electron microscopy

## Declarations

### Ethics approval and consent to participate

“Not applicable” --- This research focuses on invertebrate insects. There are no ethical considerations mentioned for these species according to EU Directive 86/609-STE123.

### Consent for publication

“Not applicable”

### Availability of data and material

The data sets supporting the results of this article will be made available in the DRYAD repository.

### Authors’ contributions

ML and VCO desiged the experiments, AA collected fly specimen, AA, SP and ML recorded fly copulation, AA performed light microscopy analysis of *Drosophila* male genitalia, FTR performed SEM analysis of *D. pachea* male genitalia. ML, AA and SP analyzed the movie datasets, ML and VCO wrote the manuscript with AA.

## Acknowledgements

We are grateful to Sadjo Sidikou and Pierre Quevreux for movie recordings, to Bertie Joan van Heuven and Menno Schilthuizen for assistance and interpretation of SEM analysis and to Martina Manns for discussions of *D. nannoptera* mating positions. We greatly thank Álvaro Barragán, Violeta Rafael and Santiago Palacios for their help in the field work made in Ecuador (Research Permit No.003-15 IC-FAU-DNB/MA), Therese Markow and Alejandro Oceguera-Figueroa for their help with fly collections in Mexico, Jean David for *Drosophila* laboratory strains and Alexandre Peluffo for help with the statistical analysis.

## Supporting Information

**Supplementary Datafile 1:** Length measurements at the left and right sides of the ventral aedeagus tip of *D. acanthoptera*, *D. nannoptera*, *D. machalilla* and *D. bromeliae*

**Supplementary Datafile 2:** Multilocus DNA sequence dataset for the molecular phylogeny shown in supplementary Fig. 6.

**Supplementary Datafile 3:** Multi-species analysis of courtship and copulation periods, shown in supplementary Fig. 6.

**Supplementary Datafile 4:** Landmark position measurements, used to calculate angle values for the multi-species mating position analysis, shown in Fig. 3.

**Supplementary Datafile 5:** Randomization of experiment names for the multi-species mating position analysis. Original names and random number substitutes are listed for each movie.

**Supplementary Datafile 6:** Copulation times of couples filmed for the position analysis of *D. nannoptera* from a frontal perspective, shown in supplementary Fig. 8.

**Supplementary Datafile 7:** Angle measurements for the position analysis of *D. nannoptera* from a frontal perspective, shown in Fig. 3.

**Supplementary Figure 1: The aedeagus of *D. pachea* is asymmetric.** Preparations in ventral view **(a)** The red circle indicates the right-sided position of the gonopore. **(a-j)** Ten preparations. The scale bar is 100 μm.

**Supplementary Figure 2: The aedeagus of *D. acanthoptera* is asymmetric.** Preparations in ventral view. **(a)** The red lines indicate the length measurements of ventral apex spurs (see materials and methods). **(a-j)** preparations. **(k)** Length measurements of apical spurs. The dashed line corresponds to the 1:1 length ratio of left and right spurs. The scale bar is 100 μm.

**Supplementary Figure 3: No asymmetry is detected in the aedeagus of *D. nannoptera*.** Preparations in ventral view **(a)** The red lines indicate the length measurements of ventral apex spurs (see materials and methods). **(a-o)** Fifteen preparations. **(p)** Length measurements of apical spurs. The dashed line corresponds to the 1:1 length ratio of left and right spurs. The scale bar is 100 μm.

**Supplementary Figure 4: No asymmetry is detected in the aedeagus of *D. machalilla*.** Preparations in ventral view **(a)** The red lines indicate the length measurements of ventral apical hooks (see materials and methods). **(a-j)** Ten preparations. **(k)** Length measurements of apical hooks. The dashed line corresponds to the 1:1 length ratio of left and right hooks. The scale bar is 100 μm.

**Supplementary Figure 5: No asymmetry is detected in the aedeagus of *D. bromeliae*.** Preparations in ventral view **(a)** The red lines indicate the length measurements of ventral apex ridges (see materials and methods). **(a-j)** Replicate preparations. **(k)** Length measurements of apex ridges. The dashed line corresponds to the 1:1 length ratio of left and right ridges. The scale bar is 100 μm.

**Supplementary Figure 6: Courtship and copulation duration in *D. pachea* and related species.** The phylogenetic relationships of analyzed species are indicated on the left with a bayesian phylogeny based on a multilocus dataset of Lang et al. 2014 [28], and additional data for *D. willistoni* and *D. tripunctata* (see material and methods, supplementary Table 1). Numbers indicate posterior probabilities for node supports < 1. Each line represents an experiment. Courtship is indicated in red, initial copulation with variable positions in blue and copulation after the settling time point in grey. Experiments are aligned by the settling time point. Time points at which the mating angle was calculated are indicated as tick marks: the settling time point in yellow; 10% stable copulation time point in green and measurements at later regular time intervals in black.

**Supplementary Figure 7: Multi-species mating position measurements.** A) Mating couple of *D. buzzatii* the scale bar 500 μm. Landmarks P1 (1), P2 (2), and P3 (3) are indicated. The dashed white lines (P1,P2) and (P2,P3) form an acute angle (semi-transparent circle sectors) which was measured to assess copulation posture. B-K) Position coordinates for angle measurements of *D. pachea* and nine related *Drosophila* species; aca: *D. acanthoptera*, bro: *D. bromeliae*, buz: *D. buzzatii*, mac: *D. machalilla*, mel: *D. melanogaster*, moj: *D. mojavensis*, nan: *D. nannoptera*, pac: *D. pachea*, tri: *D. tripunctata*, wil: *D. willistoni*. Points P1 (orange circle) are placed at coordinates (0,1) and P2 (red circle) at coordinates (0,0). P3 points are shown for the settling time-point in yellow dots, for the 10% stable copulation time point in green dots and for later time points in black circles. L) Correlation of angle values calculated from two replicate measurements (n=124) at the 10% copulation time point (Pearson correlation coefficient = 0.988, df = 121, t = 69.231, p < 10e −16). The red dashed line indicates the linear regression line.

**Supplementary Figure 8: *D. nannoptera* tilts to the right side of the female abdomen.** Frontal copulation angles (black circles) are plotted over the course of copulation. Positive and negative values indicate left-sided and right-sided angles, respectively. Grey lines connect points obtained from the same copulation couple over time. The dashed line indicates an angle of zero degrees. The position analysis from a frontal perspective is indicated on the image of the copulating couple on the right. Points and numbers indicate position landmarks P4 (4), P5 (5), and P6 (6). The dashed white lines (P1,P2) and (P2,P3) form an acute angle (semi-transparent circle sector) which corresponds to the frontal copulation angle. The scale bar is 500 μm.

**Supplementary Table 1:** Species Resources.

| Species | Source | Stock number | collection locality | collection year |
| --- | --- | --- | --- | --- |
| *D. acanthoptera* | Drosophila Species Stock Center | 15090-1693.00 | Oaxaca, Mexico | 1976 |
| *D. pachea* | Drosophila Species Stock Center | 15090-1698.02 | Sonora, Mexico | 1996 |
| *D. nanoptera* | Drosophila Species Stock Center | 15090-1692.10<br>15090-1698.12 | Oaxaca/Puebla, Mexico | 1992 |
| *D. machalilla* | Andrea Acurio |  | San Jose, Ecuador | 2015 |
| *D. bromeliae* | Drosophila Species Stock Center | 15085-1682.00 | Grand Cayman Island, UK | 1985 |
| *D. buzzatii* | Jean David |  | Bahia, Brazil | 2010 |
| *D. mojavensis* | Drosophila Species Stock Center | 15081-1352.22 | Catalina Island, USA | 2002 |
| *D. tripunctata* | Drosophila Species Stock Center | 15020-2401,02 | New Orleans, USA | 1950 |
| *D. willistoni* | Jean David |  | Rio de Janeiro, Brazil | 2010 |
| *D. melanogaster* | Drosophila Species Stock Center | 14021-0231.07 | Taiwan | 1968 |

**Supplementary Table 2:**
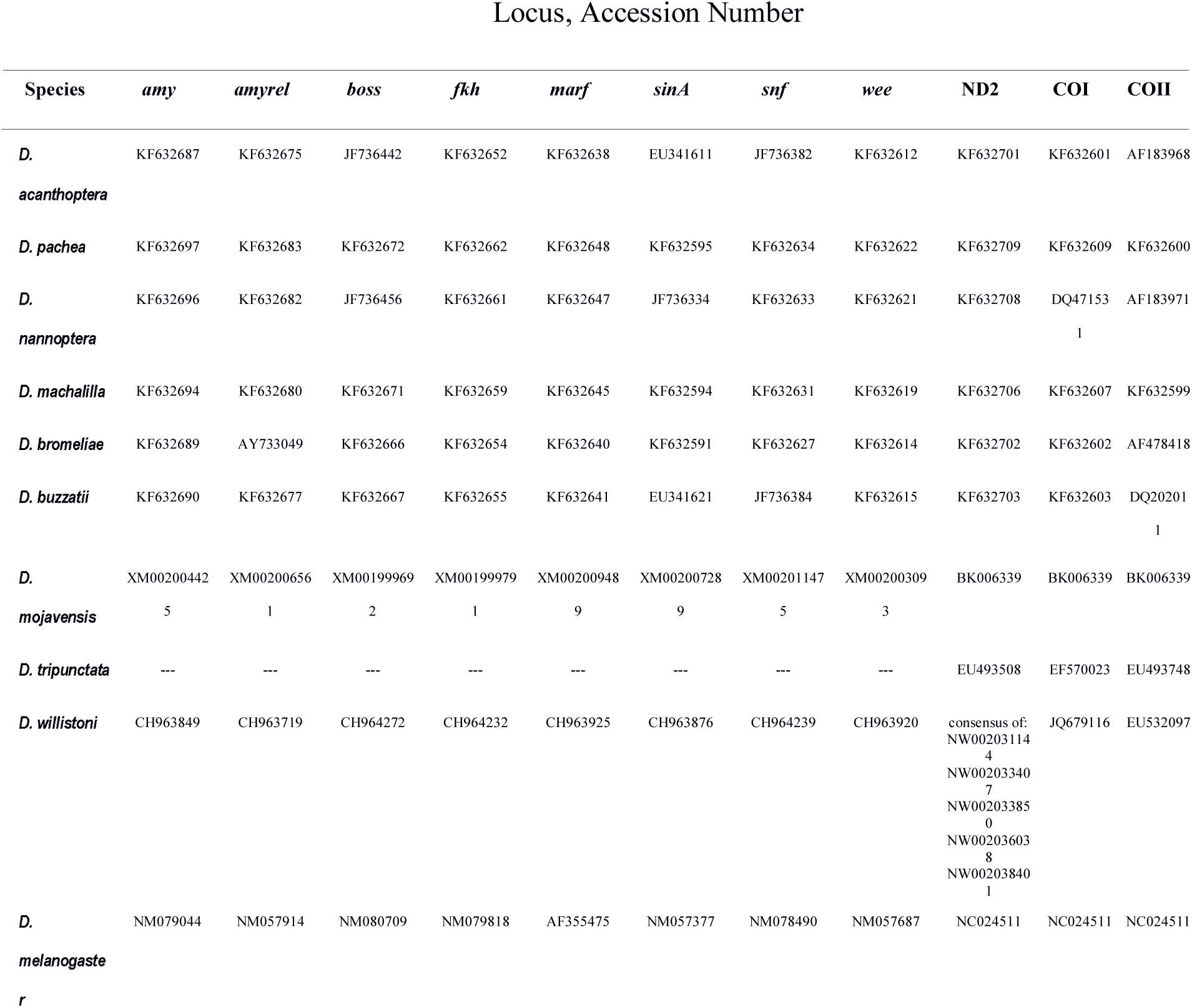
GenBank Accession Numbers of the phylogeny dataset.

