## Supplementary figures and images for "Repeated evolution of asymmetric genitalia and right-sided mating behavior in the *Drosophila nannoptera* species group"

### Supplementary Figures

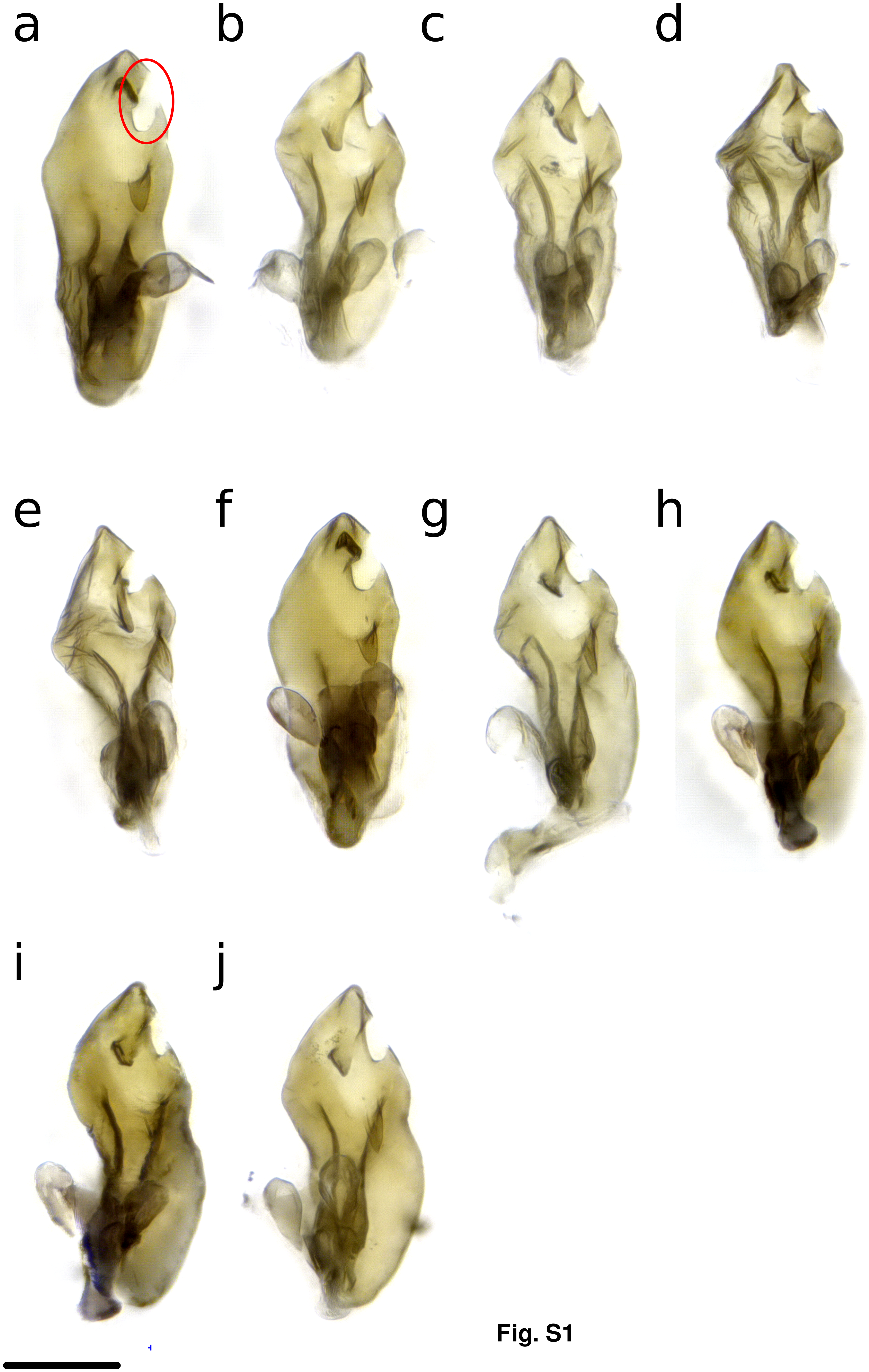


**Fig. S1**

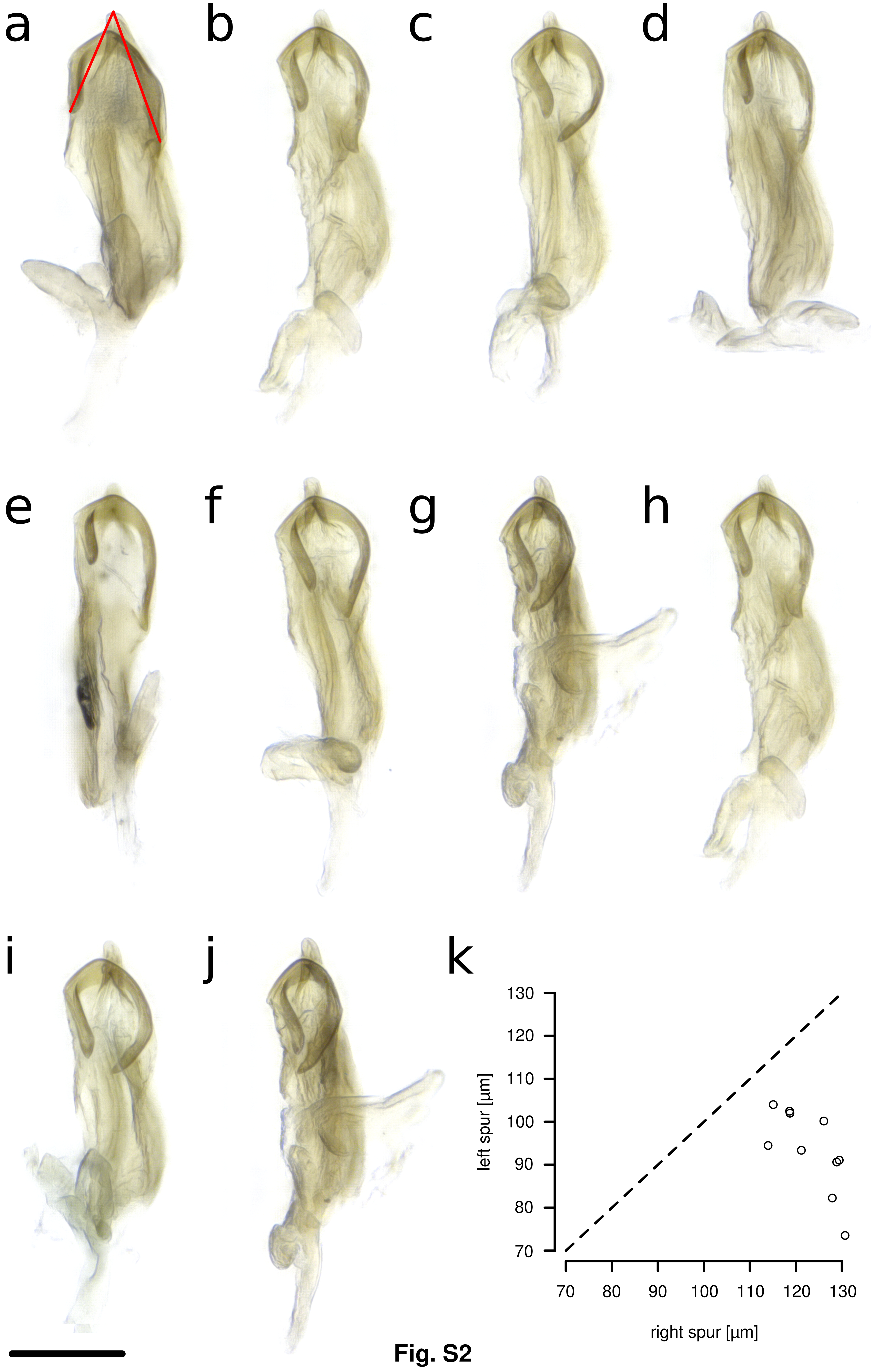

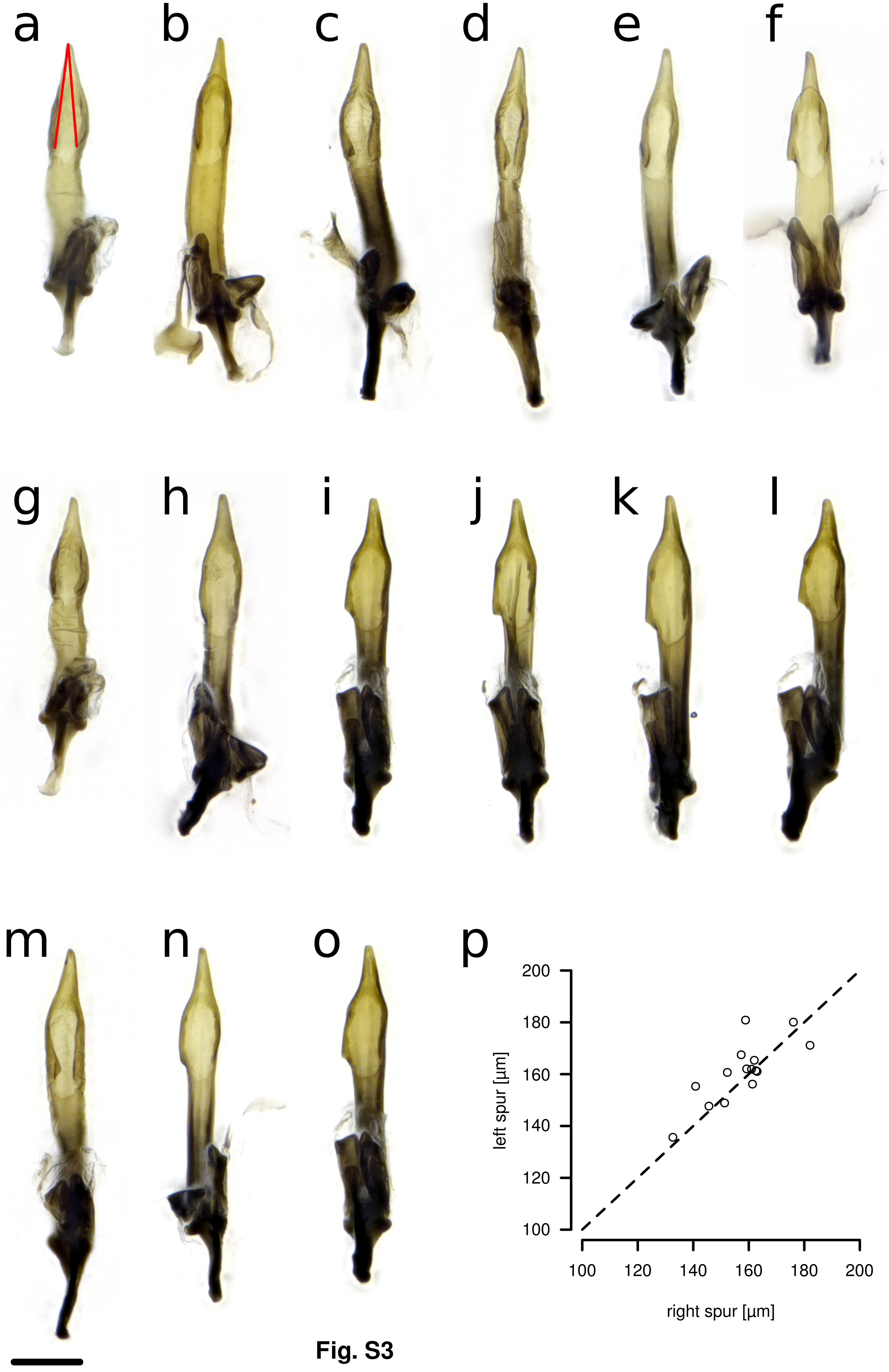

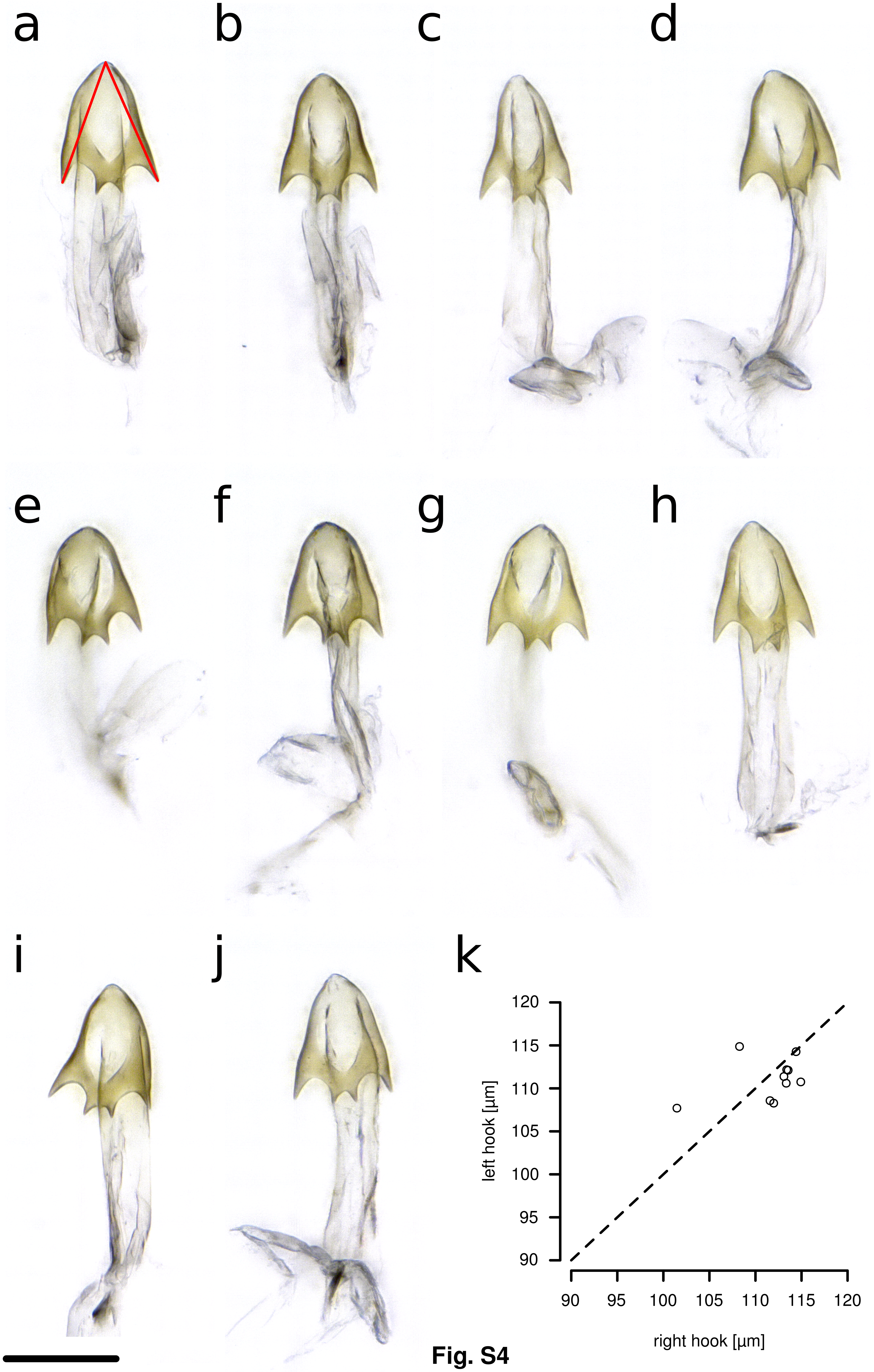

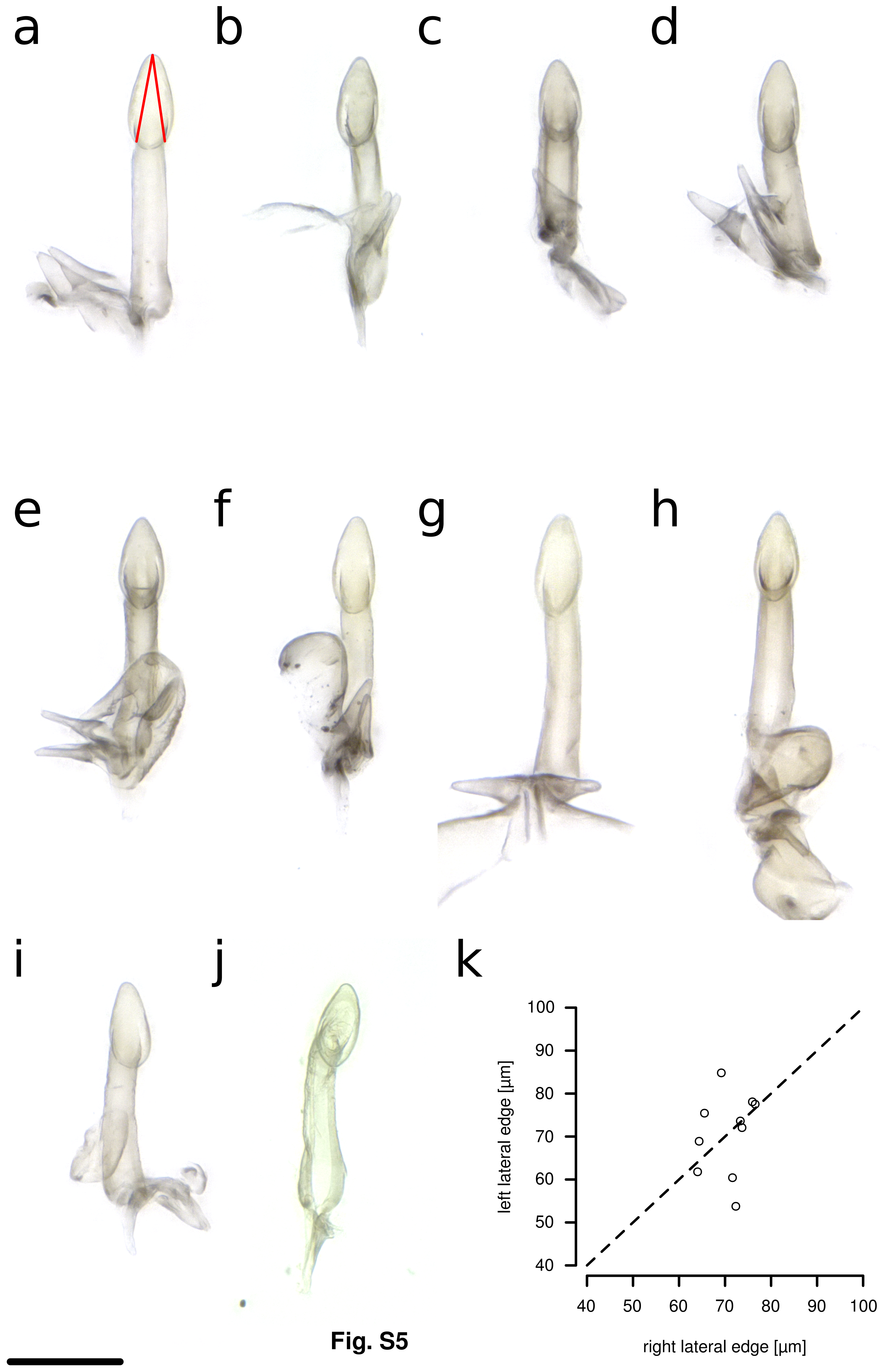

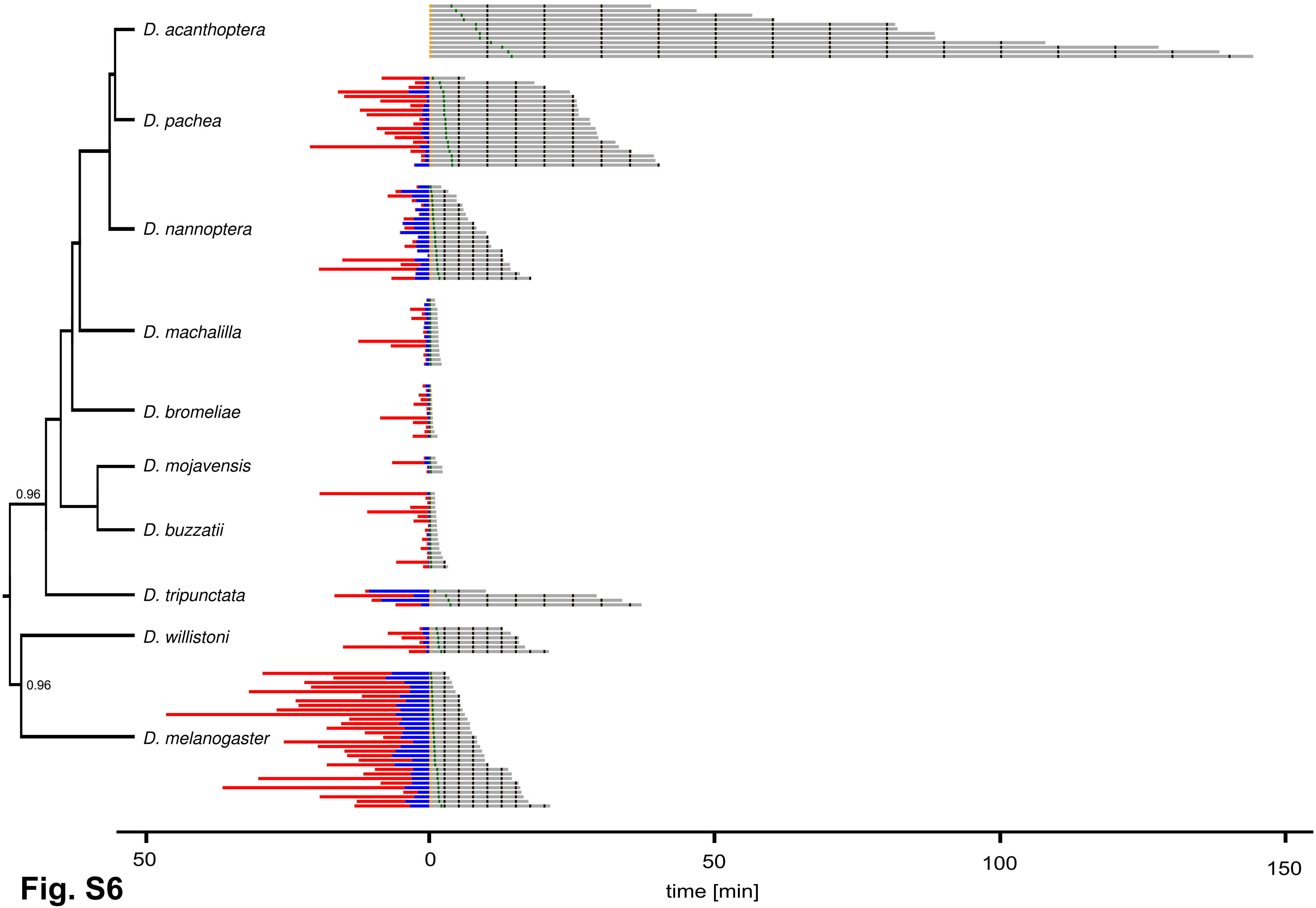

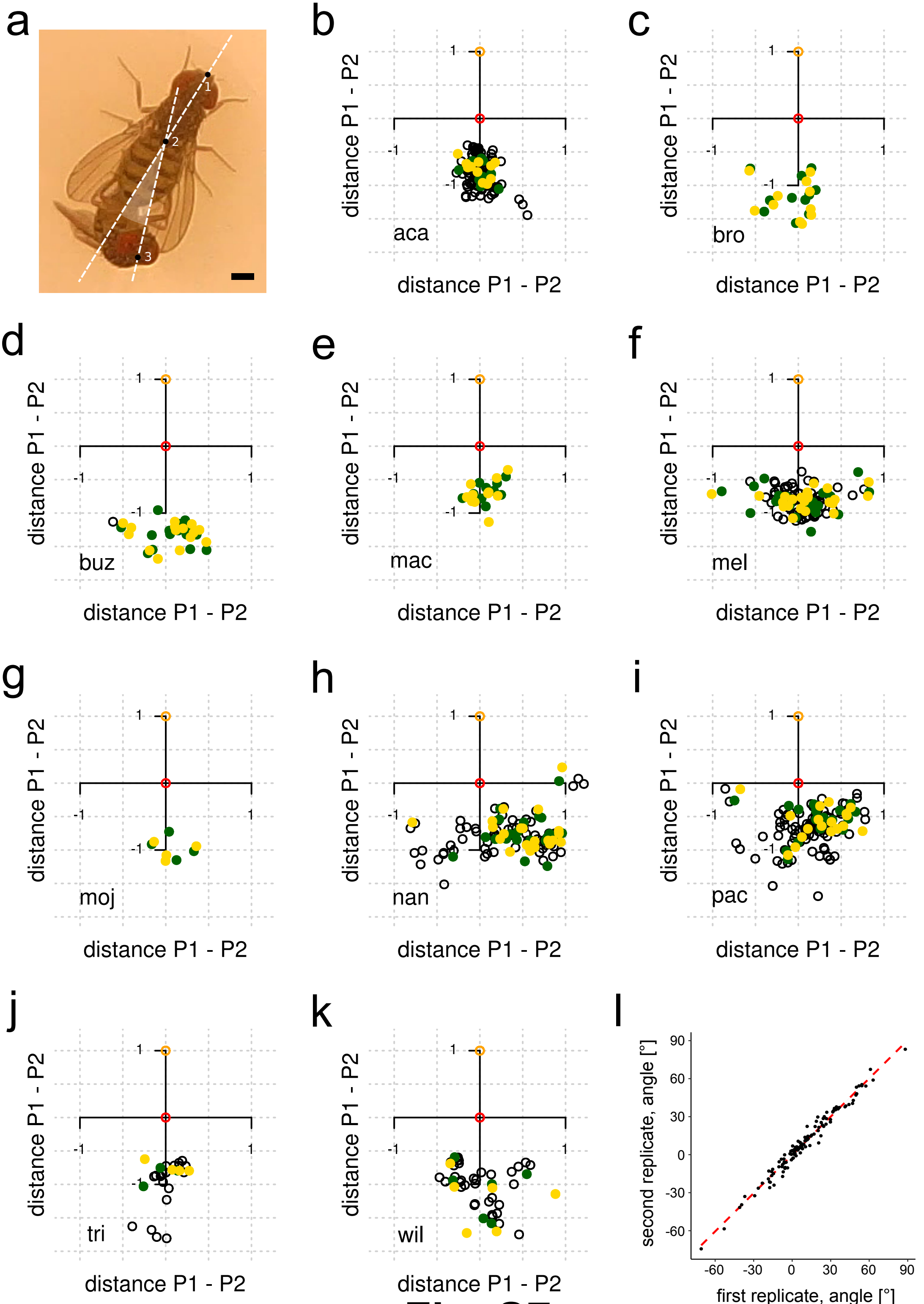

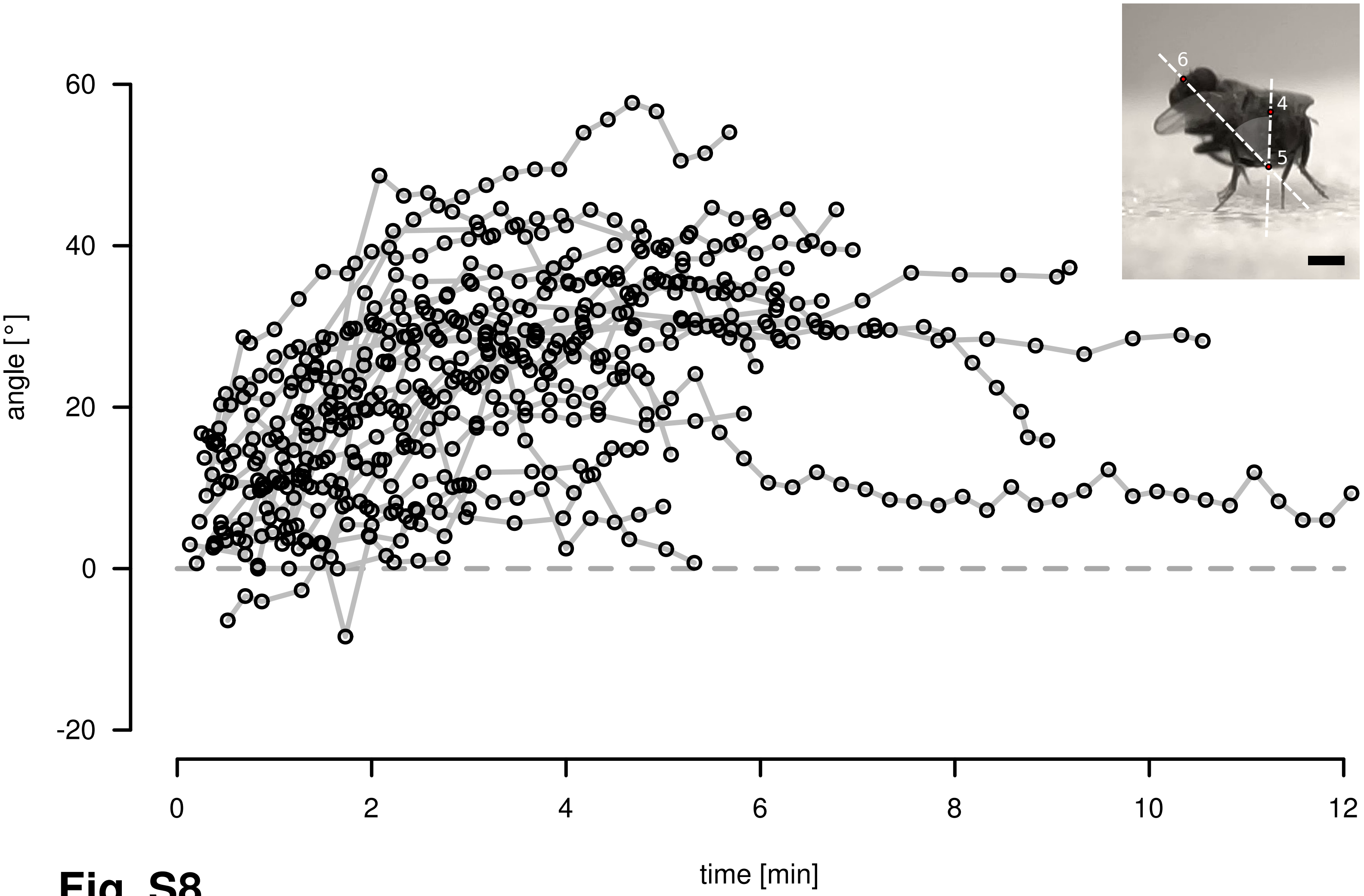


**Fig. S8**
